# The iAAA-mitochondrial protease YME1L1 regulates the degradation of the short-lived mitochondrial transporter SLC25A38

**DOI:** 10.1101/2024.05.12.593764

**Authors:** Sijie Tan, Alisa Susan Dengler, Rami Zahi Darawsheh, Nora Kory

## Abstract

Mitochondrial transporters facilitate the exchange of diverse metabolic intermediates across the inner mitochondrial membrane, ensuring an adequate supply of substrates and cofactors to support redox and biosynthetic reactions within the mitochondrial matrix. However, the regulatory mechanisms governing the abundance of these transporters, crucial for maintaining metabolic compartmentalization and mitochondrial functions, remain poorly defined. Through analysis of protein half-life data and mRNA-protein correlations, we identified SLC25A38, a mitochondrial glycine transporter, as a short-lived protein with a half-life of 4 hours under steady-state conditions. Pharmacological inhibition and genetic depletion of various cellular proteolytic systems revealed that SLC25A38 is rapidly degraded by the iAAA-mitochondrial protease YME1L1. Depolarization of the mitochondrial membrane potential induced by the mitochondrial uncoupler carbonyl cyanide m-chlorophenylhydrozone prevented the degradation of SLC25A38. This dual regulation of SLC25A38 abundance by YME1L1 and mitochondrial membrane potential suggests a link between SLC25A38 turnover, the integrity of the inner mitochondrial membrane, and electron transport chain function. These findings open avenues for investigating whether mitochondrial glycine import coordinates with mitochondrial bioenergetics.

## INTRODUCTION

Mitochondria are metabolic hubs for energy production, biosynthetic processes, and redox reactions^1^. Transport proteins, including members of the <u>S</u>o<u>L</u>ute <u>C</u>arrier 25 (SLC25) family, facilitate the transport of metabolites across the inner mitochondrial membrane (IMM) to ensure adequate concentrations of critical substrates and cofactors in the mitochondrial matrix to support these reactions^2^. However, the coordination of SLC25 transporter abundance and activity with cellular metabolic requirements, as well as the regulation of mitochondrial metabolite pools and functions, remain poorly defined.

Glycine is an important metabolic precursor for mitochondrial processes and cellular metabolism. Mitochondrial glycine metabolism is intimately linked with heme synthesis^3–5^ and one-carbon metabolism^6^. The availability of glycine in mitochondria can be controlled in part by its import through SLC25A38. Notably, the uptake of glycine into the mitochondrial matrix through SLC25A38 is required for the initiation of heme synthesis^3,4^. Loss of this transporter significantly reduces levels of 5- aminolevulinate, the first product in the heme biosynthetic pathway, as well as downstream levels of heme and mitochondrial cytochromes. In humans, mutations in *SLC25A38* cause congenital sideroblastic anemia^7^, a condition where iron accumulates in the mitochondria of erythroblasts due to dysfunctional heme synthesis or processing.

Despite the pivotal roles of glycine in mitochondrial metabolic processes and functions, the coordination of SLC25A38 abundance with mitochondrial metabolic demands remains unknown. In this study, we mined protein half-life data and mRNA-protein abundance ratios to investigate the regulation of mitochondrial transporters. We identified SLC25A38 as a short-lived mitochondrial transporter that is rapidly turned over by the iAAA-mitochondrial protease YME1L1 under physiological conditions.

## RESULTS

### SLC25A38 is a short-lived mitochondrial protein

To identify regulated mitochondrial transporters, we compared the protein and mRNA levels of SLC25 transporters using information from OpenCell^8^. While the protein-to-mRNA abundance ratios for most SLC25 transporters followed the correlation observed in the general proteome, SLC25A23, SLC25A35, SLC25A38, SLC25A39, and SLC25A42 exhibited low ratios (Fig. 1a), suggesting potential regulation of their synthesis or degradation. Indeed, SLC25A39, the mitochondrial glutathione transporter^9,10^, has recently been reported as a short-lived protein regulated by proteolysis^11,12^. Based on this, we hypothesized that SLC25A38 protein is also rapidly turned over under steady-state conditions. To test this hypothesis, we mined published proteomics datasets for half-life data on SLC25 transporters. High-throughput quantitative proteomics mapping protein degradation kinetics identified SLC25A38 as a short-lived protein in human cell lines under basal conditions^13,14^ (Fig. 1b). To measure the half-life of SLC25A38 in HEK293T cells, we blocked protein synthesis using cycloheximide (CHX) and determined SLC25A38 protein levels over time (‘chase’) by Western blot analysis (Fig. 1c). We observed that endogenous and stably expressed FLAG-tagged SLC25A38 had a half-life of ∼4 h (Fig. 1c), significantly shorter than the median half-life of several days for mitochondrial carriers in mammalian cells^13^ (Fig. 1b). Conversely, SLC25A51, a mitochondrial NAD^+^ transporter^15–17^ with similar protein-to-mRNA abundance ratios as the general proteome, exhibited a half-life of >8 h (Fig. 1c). This highlights that the turnover of SLC25A38 under basal conditions is specific to this transporter and not due to general destabilization of mitochondrial transporters by CHX. To further ascertain whether the rapid degradation of SLC25A38 was due to the removal of its mitochondrial pool, we blocked protein synthesis using CHX, fractionated cells at various time points post-treatment into cytosolic and mitochondrial fractions, and measured SLC25A38 protein levels in both fractions by Western blot. We observed that endogenous and FLAG-tagged SLC25A38 were primarily present in the mitochondrial fraction, and that their protein levels decreased over time following subsequent CHX treatment (Fig. 1d). Immunofluorescence imaging also highlighted the distinct localization of SLC25A38 within mitochondria, with the signal decreasing upon CHX treatment (Fig. 1e). These observations indicate that under physiological conditions, SLC25A38 is a short-lived mitochondrial protein due to rapid degradation.

**Figure 1:**
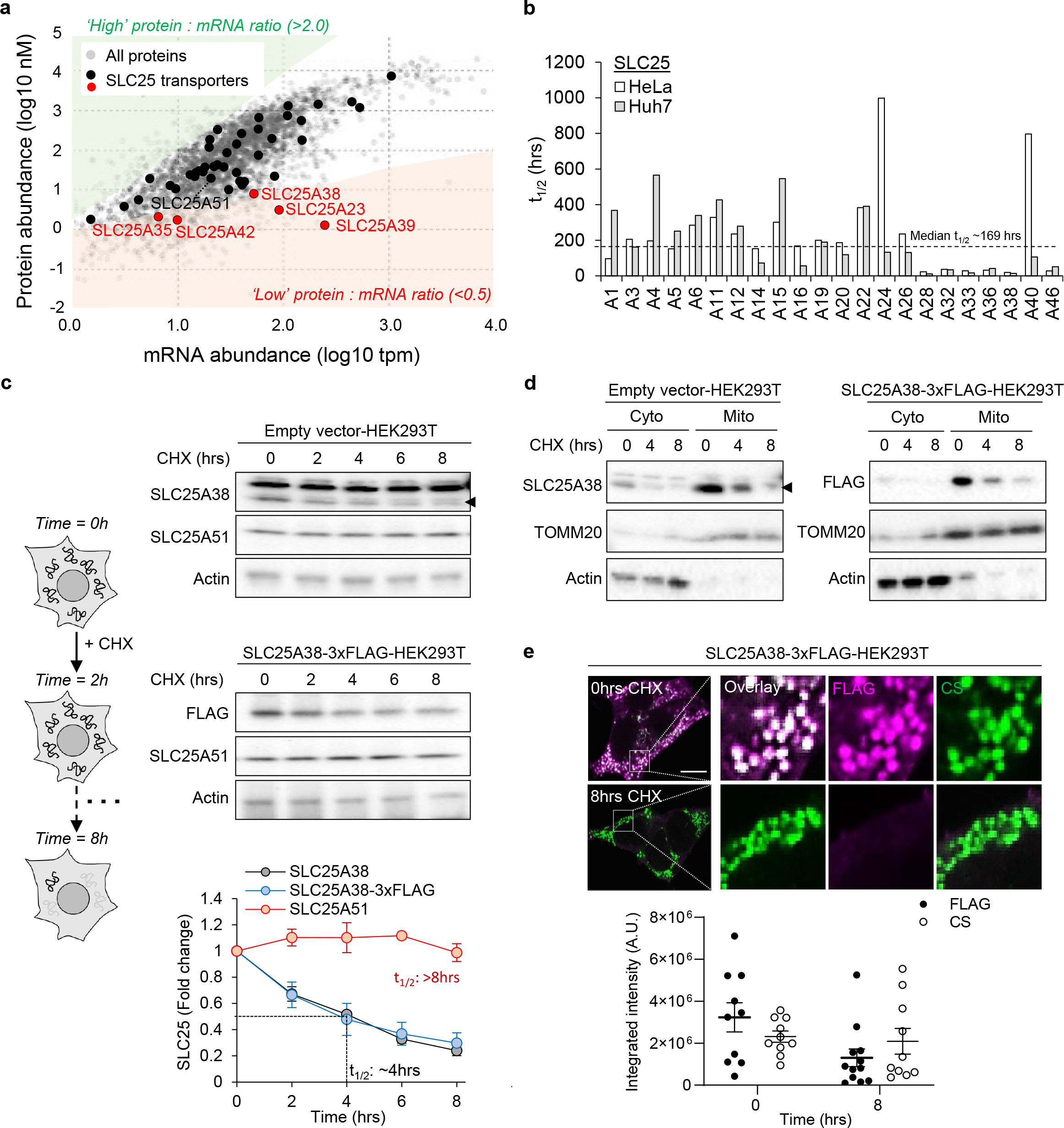
SLC25A38 is a short-lived mitochondrial transporter with a half-life of ∼4 h. (a) Scatterplot from OpenCell^8^ showing the protein and mRNA abundance of all proteins expressed in HEK293T cells. The red region highlights proteins with a low protein-to-mRNA ratio defined by a cut-off ratio of ∼0.5. The green region highlights proteins with a high protein-to-mRNA ratio defined by a cut- off ratio of 2.0. (b) The half-life of SLC25 transporters in HeLa and Huh7 cells. Graph is plotted with data extracted from the human mitochondrial high-confidence proteome (MitoCoP^13^). The dotted line indicates the median half-life of mitochondrial transporters. (c) Relative protein abundance of endogenous and FLAG-tagged SLC25A38, and endogenous SLC25A51 at different CHX treatment time points normalized to the protein abundance at 0 h (n = 3). For SLC25A51, the levels in SLC25A38- 3xFLAG-HEK293T were plotted. (d) Cytosolic and mitochondrial levels of endogenous and FLAG-tagged SLC25A38 at different CHX treatment time points. TOMM20 and actin served as controls to indicate the purity of cellular fractionation (n = 1). (e) Quantification of FLAG and citrate synthase (CS) signals in SLC25A38-3xFLAG-HEK293T cells under 0 and 8 h of CHX treatment. A total of 10 images with 200–400 cells for each time point were quantified to calculate the average FLAG and CS intensity. Scale bar = 10 μm. Values are mean ± S.D.M.

### Proteasomal and lysosomal systems do not regulate the stability or turnover of SLC25A38

We sought to determine the machinery involved in SLC25A38 degradation, focusing on the three major proteolytic systems that contribute to the degradation of mitochondrial proteins: (i) the proteasome, (ii) lysosomes (mitophagy), and (iii) mitochondrial proteases. Inhibiting the proteasome using bortezomib and epoxomicin did not impact the basal levels or stability of SLC25A38 (Fig. 2a). Likewise, treatment with NH4Cl to inhibit lysosomal acidification or using *Atg5* knockout (KO) to block autophagy did not alter the basal levels or turnover rate of SLC25A38 (Fig. 2b and c). These results highlight that the degradation of SLC25A38 occurs independently of the proteasome and mitophagy.

**Figure 2:**
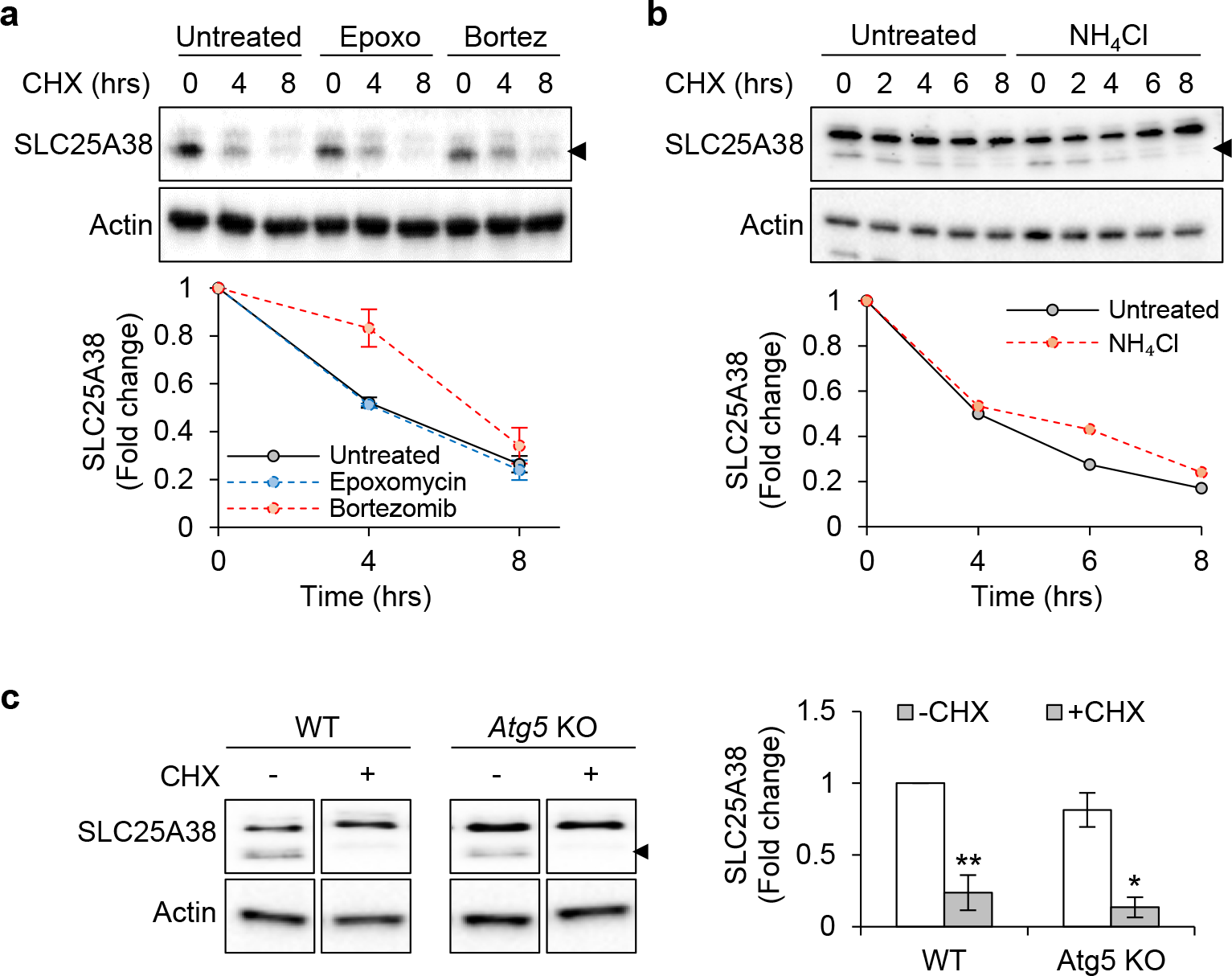
Proteasomal inhibition, lysosomal inhibition, and *Atg5* KO do not affect SLC25A38 degradation under basal conditions. (a–b) Relative SLC25A38 levels at different CHX treatment time points under basal conditions (untreated), proteasomal inhibition with bortezomib or epoxomycin, or lysosomal inhibition with NH4Cl. Fold change values for SLC25A38 under the different inhibitions were normalized to the protein abundance at 0 h (n = 2 for a, n = 1 for b). (c) Relative SLC25A38 levels in WT and *Atg5* KO HEK293T cells with and without CHX treatment for 16 h (n = 3). Cropped blots were from the same gel. Fold change values for SLC25A38 in *WT* and *Atg5* KO were normalized to WT - CHX. Values are mean ± S.D.M.

### The mitochondrial iAAA protease YME1L1 degrades SLC25A38 under basal conditions

As an IMM protein, SLC25A38 is accessible to proteases present in the mitochondrial matrix, inner membrane, and intermembrane space, which include at least 20 catalytically functional proteases involved in cellular stress signaling and proteostasis^18,19^. To investigate the contribution of mitochondrial proteases to SLC25A38 turnover, we used CRISPR to delete five major mammalian mitochondrial proteases and monitored SLC25A38 degradation using a CHX-chase assay. Deletion of AFG3L2, LONP1, CLPP, and PARL did not affect SLC25A38 levels or turnover (Fig. 3a). However, YME1L1 KO increased the steady-state levels of SLC25A38 and substantially extended the transporter’s half-life to over 8 h (Fig. 3b). Immunoprecipitation of FLAG-tagged SLC25A38 revealed YME1L1 as a binding partner under basal conditions (Supplementary Fig. 1). These results suggest that YME1L1 regulates the rapid degradation of SLC25A38 basally.

**Figure 3:**
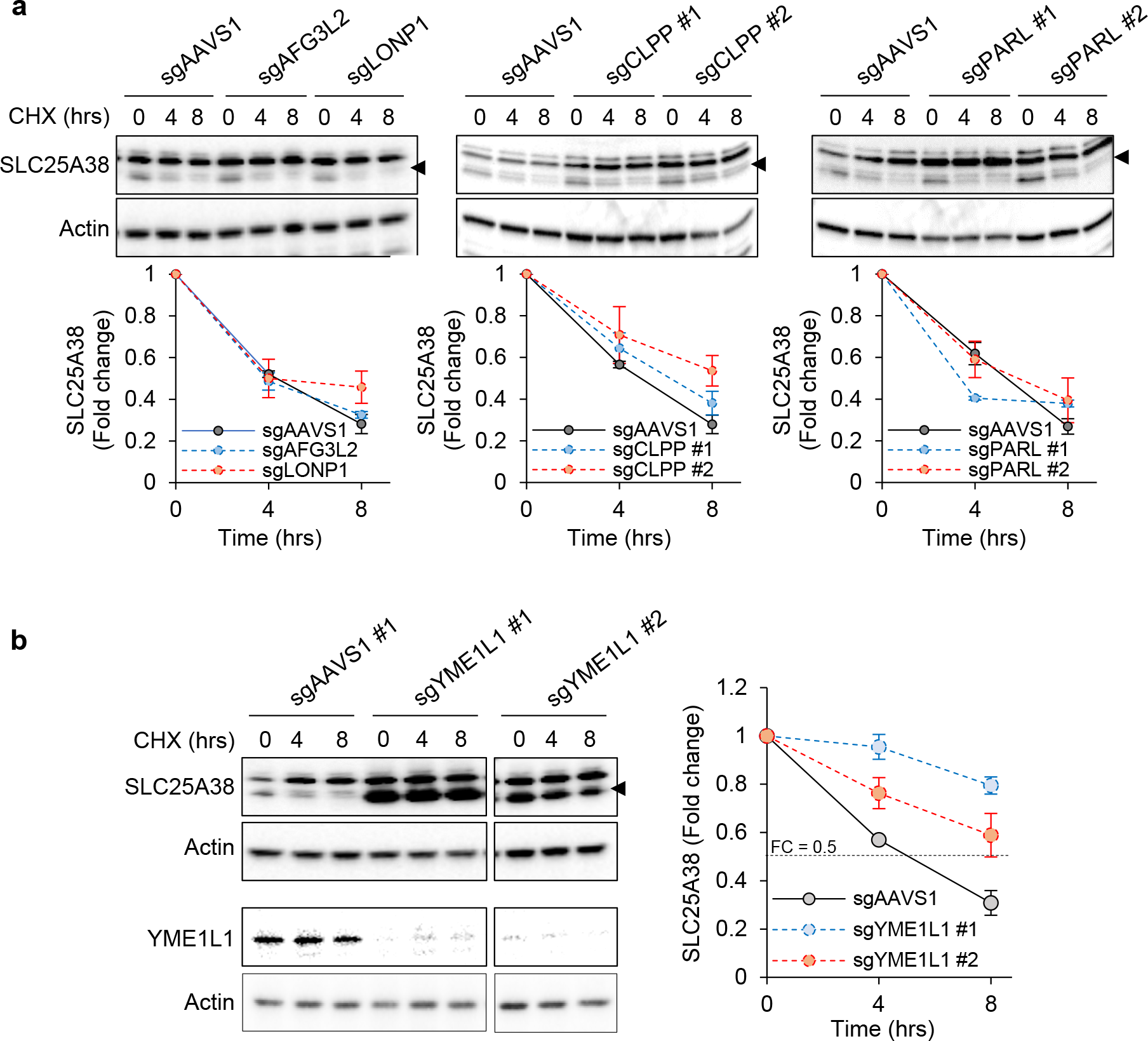
YME1L1 regulates SLC25A38 degradation under basal conditions. (a) Immunoblots and relative SLC25A38 levels in non-targeting KO control (sgAAVS1) and mitochondrial protease CRISPR KO cells at different time points of CHX treatment (n = 2). (b) Immunoblots and relative SLC25A38 levels at different CHX treatment time points in non-targeting sgAAVS1 control and two YME1L1 KO clones (n = 3). Immunoblot for YME1L1 shows the depletion of protease in the KO clones. Fold-change values for SLC25A38 in the different mitochondrial protease KO cells were normalized to the protein abundance at 0 h. Values are mean ± S.D.M.

### CCCP treatment prevents SLC25A38 degradation

Next, we explored the cellular cues that could regulate SLC25A38 stability. Given that the first step of heme synthesis requires the uptake of glycine into the mitochondrial matrix via SLC25A38^3,4^, we investigated whether glycine starvation or perturbation of heme biosynthesis affects the stability of SLC25A38. As glycine can be synthesized from serine in the mitochondrial matrix, we evaluated SLC25A38 degradation in media depleted of glycine and/or serine. However, neither serine nor glycine starvation affected SLC25A38 stability (Supplementary Fig. 2a). Similarly, inhibition of heme synthesis with succinylacetone (Supplementary Fig. 2b) or the addition of hemin (Supplementary Fig. 2c) did not affect SLC25A38 degradation. Since the rate-limiting enzyme of heme biosynthesis, 5-aminolevulinate (ALA) synthase, requires pyridoxial-5’-phosphate (PLP) as a cofactor for its activity^5^, we also cultured HEK293T cells in PLP-depleted media supplemented with low (1 nM) or excess (1 μM) pyridoxine, the metabolic precursor of PLP, to monitor SLC25A38 stability. However, perturbation of pyridoxine levels did not affect SLC25A38 degradation (Supplementary Fig. 2d). These results suggest that the availability of metabolites involved in heme biosynthesis does not influence SLC25A38 turnover.

As YME1L1 is a stress-sensitive mitochondrial protease responsive to oxidative stress, hypoxia, starvation, and mitochondrial membrane depolarization^20,21^, we investigated whether similar cellular and mitochondrial stressors affect SLC25A38 stability. H2O2-induced oxidative stress, serum starvation, and inhibition of electron transport chain (ETC) complex with antimycin A (a complex III inhibitor) or oligomycin (an ATP synthase inhibitor) did not affect SLC25A38 degradation (Fig. 4a). However, the addition of the mitochondrial uncoupler CCCP prevented SLC25A38 degradation and extended the transporter half-life beyond 8 h (Fig. 4a). Interestingly, CCCP treatment did not affect the levels of YME1L1 (Fig. 4b).

**Figure 4:**
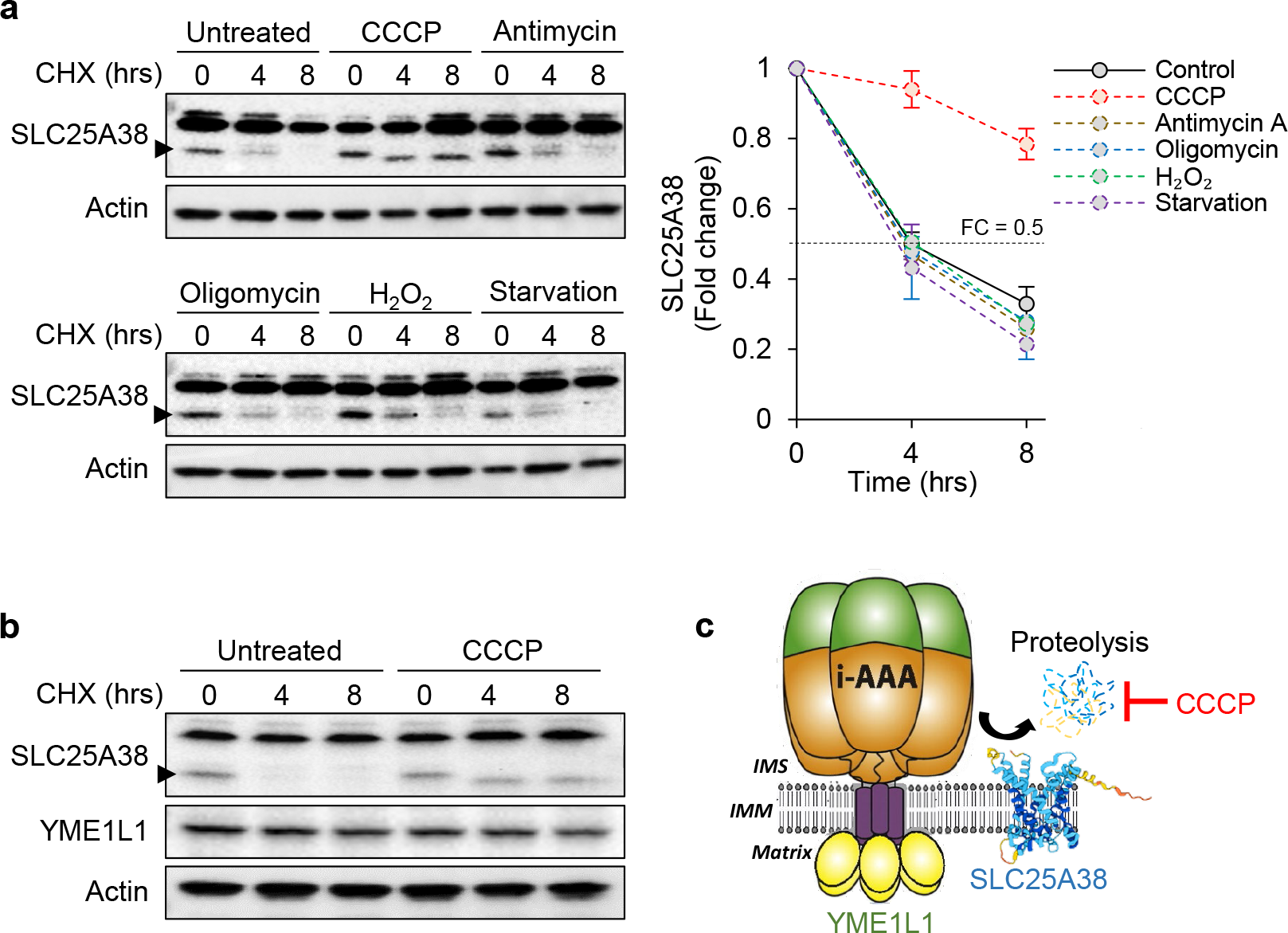
CCCP prevents SLC35A38 degradation. (a) Immunoblots and relative SLC25A38 levels at different CHX treatment time points under basal (untreated), CCCP, antimycin A, oligomycin, H2O2, and serum starvation conditions. Fold-change values for SLC25A38 under the different stress conditions were normalized to the protein abundance at 0 h (n = 2–3). Values are mean ± S.D.M. (b) Representative immunoblot for YME1L1 levels under untreated or CCCP treatment conditions, in the presence of CHX at different time points. (c) Schematic diagram for the dual regulation of SLC25A38 stability by YME1L1 and CCCP.

## DISCUSSION

SLC25 transporters comprise the largest solute carrier family in humans, facilitating the import and export of diverse solutes across the IMM^2^. They function as gatekeepers to compartmentalize and protect intramitochondrial metabolite pools, creating an optimal chemical environment to support numerous biochemical processes within the matrix. Regulating the levels of mitochondrial transporters could provide a mechanism to control the content and activity of mitochondrial metabolism^1^. However, the regulatory mechanisms that finely tune transporter abundance to coordinate metabolic homeostasis remain largely unknown for most SLC25 transporters. Here, we report that the mitochondrial glycine transporter SLC25A38^3,4^ is a short-lived protein whose abundance is regulated by the iAAA- mitochondrial protease YME1L1. By coupling a CHX-chase assay with pharmacological inhibition or genetic depletion of various cellular proteolysis machinery, our study confirmed that the stability and turnover rate of SLC25A38 depend on YME1L1 under basal conditions. This aligns with a previously published proteomics dataset showing that SLC25A38 accumulates in YME1L1 KO cells, along with other YME1L1 substrates^21^. Thus, together with other recent studies^11,12^, our research establishes degradation by mitochondrial proteases as a mechanism regulating the abundance of transporters in the IMM. While our results indicate a direct interaction between YME1L1 and SLC25A38, further experiments are required to determine whether YME1L1 directly degrades SLC25A38, or if YME1L1 indirectly influences SLC25A38 stability through the degradation of other proteases^22,23^, as reported for OMA1.

Glycine participates in multiple metabolic processes in the mitochondria. Notably, the first step of heme synthesis requires glycine import into the mitochondrial matrix through SLC25A38^3,4^. Gene ontology annotations associate SLC25A38 with erythrocyte differentiation, and loss-of-function mutations in SLC25A38 lead to congenital sideroblastic anemia^24^. Given that glycine import is the rate-limiting step in heme synthesis, we expected that changes in glycine and heme availability would trigger a compensatory mechanism to fine-tune SLC25A38 abundance on the IMM, analogous to the proteolytic regulation of SLC25A39 by mitochondrial glutathione levels^11,12^. However, perturbation of heme biosynthesis and levels —such as inhibition of ALA dehydratase, cofactor PLP starvation, or addition of hemin—did not affect SLC25A38 turnover. Similarly, modulation of intracellular serine and/or glycine pools via metabolite starvation did not affect SLC25A38 stability. As these experiments were conducted in non-erythroid HEK293T cells, additional experiments using erythroid cell lines such as K562 or HEL will be pivotal to confirm the effects of impaired heme biosynthesis on SLC25A38 stability.

While glycine and heme availability did not affect SLC25A38 turnover, we observed an unexpected stabilization of SLC25A38 upon mitochondrial uncoupling with CCCP. Hem25, along with FLX1 (the yeast homologs of human SLC25A38 and SLC25A32, respectively), have been reported to play important roles in maintaining the integrity of respiratory complexes and ETC activity^25^. Loss of Hem25 and FLX1 destabilized specific subunits from complex I to III, likely due to reduced heme levels, which is an essential component of ETC complexes^25^. We postulate that disruption of the mitochondrial membrane proton gradient and ETC activity by CCCP triggers a feedback mechanism to alleviate ETC impairment via SLC25A38 stabilization. As we did not observe an effect of antimycin A on SLC25A38 turnover, we suspect that the specific inhibition of complex III subunits may inadequately induce ETC impairment to affect SLC25A38 stability. In addition, the absence of a heme moiety in ATP synthase might explain the lack of effect on SLC25A38 stability by oligomycin. Yme1L1 levels have previously been reported to be regulated by membrane potential^20^. Interestingly, we did not observe a change in YME1L1 levels with CCCP treatment. The mechanism by which SLC25A38 is stabilized upon CCCP treatment requires investigation. Understanding the regulatory mechanisms of SLC25A38 under physiological conditions or during ETC impairment may unveil novel roles SLC25A38 in mitochondrial bioenergetic processes.

## MATERIALS AND METHODS

### Cell culture

HEK293T cells were cultured in DMEM high glucose with L-glutamine and sodium pyruvate supplemented with 10% fetal bovine serum (FBS) and 1% penicillin–streptomycin (Hyclone, Cytiva) (‘regular media’), unless stated otherwise. All cell lines were grown in 5% CO2 at 37°C.

### Lentivirus production and generation of stable cell lines

For lentivirus production, 400,000 HEK293T cells were seeded in a 6-well plate in DMEM with 20% heat-inactivated FBS (IFS) and 1% penicillin–streptomycin. After a 24-h recovery, the cells were transfected with 1 μg pMXs empty vector or 1 μg pMXs-SLC25A38-3xFLAG, 900 ng pCMV-Gag-Pol, and 150 ng pCMV-VSVG using X-tremeGENE 9 DNA Transfection Reagent (Sigma) following the manufacturer’s protocol. At 24 h later, the media was replaced with DMEM containing 30% IFS and 1% penicillin–streptomycin. Another 24 h later, the supernatant was collected and filtered through a 0.45 μm sterile filter. The virus was then stored at -80°C. For infection, 400,000 HEK293T cells were seeded in a 6-well plate. After a 24-hour recovery, the cells were infected with 300 μl of virus and 10 μg/ml polybrene (Millipore). After a 48-hr infection, the cells were selected with 10 μg/ml blasticidin (Millipore) until the control well (without virus addition) reached 100% cell death. The empty vector-HEK293T and SLC25A38-3xFLAG-HEK293T stable cells were subsequently maintained in 2 μg/ml blasticidin.

### CHX-chase assay

For the CHX-chase assay, 800,000 cells were seeded in a 6-well plate. After a 24-h recovery, cells were treated with 10 μg/ml CHX (Sigma) in regular media (untreated), or with the addition of 10 nM bortezomib (MedChemExpress), 50 nM epoxomicin (MedChemExpress), 20 mM NH4Cl (Sigma), 50 μM CCCP (Sigma), 1μM antimycin A (Sigma), 1μM oligomycin (Sigma), or 200 μM H2O2 (Sigma) and harvested at the indicated time points. For serum starvation, cells were treated with 10 μg/ml CHX in regular DMEM without the addition of FBS.

### Serine, glycine, and pyridoxial-5’-phosphate starvation

For serine and/or glycine depletion experiments, 500,000 cells were seeded in a 6-well plate with regular media. After a 24-hour recovery, the regular media was removed and the cells were further cultured in DMEM/F12 media without amino acids and glucose (US Biologicals), supplemented with 10% double-dialyzed FBS, 1% penicillin–streptomycin, 25 mM glucose, 1x essential amino acids mix, 1x non-essential amino acids mix without serine and glycine, as well as 0.4 mM serine and/or glycine as per experimental needs. After 24 h, cells were further treated with 10 μg/ml CHX in the respective media.

For PLP starvation experiments, 200,000 cells were seeded in a 6-well plate with regular media. After a 24-h recovery, the regular media was removed and the cells were further cultured in RPMI 1640 media without L-glutamine, vitamins, and glucose (US Biologicals), supplemented with 5% hydroxylamine-treated dialyzed IFS^26^, 10 mM glucose, 1% penicillin–streptomycin, a vitamin mix lacking vitamin B6 prepared in-house following the concentrations in the 1x RPMI vitamin mix (Sigma), and 1 nM or 1 μM pyridoxine (Sigma) as per experimental needs. After 72 h, the cells were further treated with 10 μg/ml CHX in the respective media.

### Perturbation of intracellular heme levels

A total of 500,000 cells were seeded in a 6-well plate. After a 24-h recovery, cells were further cultured in regular media (control), or regular media supplemented with 0.5 mM succinylacetone (MedChemExpress) or 25 μM hemin (MedChemExpress). After 24 h, cells were further treated with 10 μg/ml CHX in the respective media.

### Protein lysis and immunoblotting

Cells were lysed in RIPA buffer (Thermo Scientific) containing 1x Halt Protease and Phosphatase Inhibitor Cocktail (Thermo Scientific). The cells were lysed on ice for 20 min with intermittent vortexing, followed by centrifugation at 14,000 rpm for 15 min at 4°C. Supernatants were collected. Cellular fractionation into cytosolic and mitochondrial fractions was performed using the Mitochondria Isolation Kit for Cultured Cells (Thermo Scientific) following the manufacturer’s protocol. Protein concentrations were determined with the Pierce BCA Protein Assay Kit (Thermo Scientific). SDS-PAGE and wet transfer Western blotting were performed, and the blots were labeled with Pierce™ ECL Western Blotting Substrate (Thermo Scientific) and imaged using the Bio-Rad ChemiDoc MP imaging system.

The following antibodies and dilutions were used: β-actin (13E5) HRP conjugate (Cell Signaling #5125; 1:10,000), FLAG (M2) (Sigma F1804; 1:1,000), SLC25A38 (Abcam ab133614; 1:1,000), SLC25A51 (Novus NBP2-46847; 1:1,000), TOMM20 (Proteintech 11802-1-AP; 1:1,000), anti-rabbit IgG HRP-linked antibody (Cell Signaling 7074; 1:10,000), and anti-mouse IgG HRP-linked antibody (Cell Signaling 7076; 1:10,000).

### Co-immunoprecipitation

Approximately 10 million HEK293T cells stably expressing either the empty vector or SLC25A38- 3XFLAG were lysed on ice in 500 μl co-IP buffer (50 mM Tris base pH 7.5, 150 mM NaCl, 1mM EDTA, 1% triton X-100) containing 1x Halt Protease and Phosphatase Inhibitor Cocktail for 20 min, followed by centrifugation at 14,000 rpm for 15 min at 4°C to collect the protein lysates. A total of 15 μl of protein lysates was collected as ‘input’, and the remaining were transferred to Eppendorf tubes containing 40 μl of pre-cleared FLAG-M2 magnetic beads (Sigma). The protein lysates and beads were mixed and incubated at 4°C overnight. The next day, the beads were washed with buffer (50 mM Tris base pH 7.5, 150 mM NaCl), and FLAG-bound proteins were eluted with 4x Laemmli sample buffer (Bio-Rad). The samples were boiled for 3 min before undergoing SDS-PAGE.

### Immunofluorescence

Approximately 200,000 cells were seeded onto poly-D-lysine-coated coverslips (Neuvitro). After a 24-h recovery, the cells were treated with 10 μg/ml CHX and fixed with 3% paraformaldehyde (Sigma) and 0.1% glutaraldehyde (Sigma) for 10 min at the indicated time points. Subsequently, the cells were permeabilized and blocked using a buffer containing 0.2% milk, 2% FBS, 1% BSA, and 0.01% Triton X-100. The cells were stained with FLAG and citrate synthase (D7V8B) (Cell Signaling #14309) antibodies (1:300 dilution) overnight, followed by Abberior STAR secondary antibodies (1:800 dilution) for 2 h. Coverslips were mounted onto slides with Prolonged Gold Antifade Medium containing DAPI (Invitrogen). Images were acquired with Zeiss spinning disk confocal microscope using the 100x oil immersion objective lens. A total of 10 random fields per coverslip were imaged, and the same acquisition parameters were applied to all conditions.

### Cloning single-guide RNA (sgRNA) into pX330

sgRNAs were designed using the Synthego Knockout Guide Designer tool. The sgRNA sequences are reported in Table 1. Forward and reverse sgRNA oligos were annealed at 95°C for 5 min, followed by slow cooling to 4°C. pX330 (Addgene) was digested with *BbsI*. The linearized pX330 was separated on a DNA agarose gel, and the band was excised and purified with the QIAquick Gel Extraction Kit (Qiagen). Annealed oligos were ligated with linearized pX330 at room temperature overnight with T4 ligase (NEB). Ligated products were transformed into DH5α, and positive clones were confirmed by Sanger sequencing.

**Table 1:**
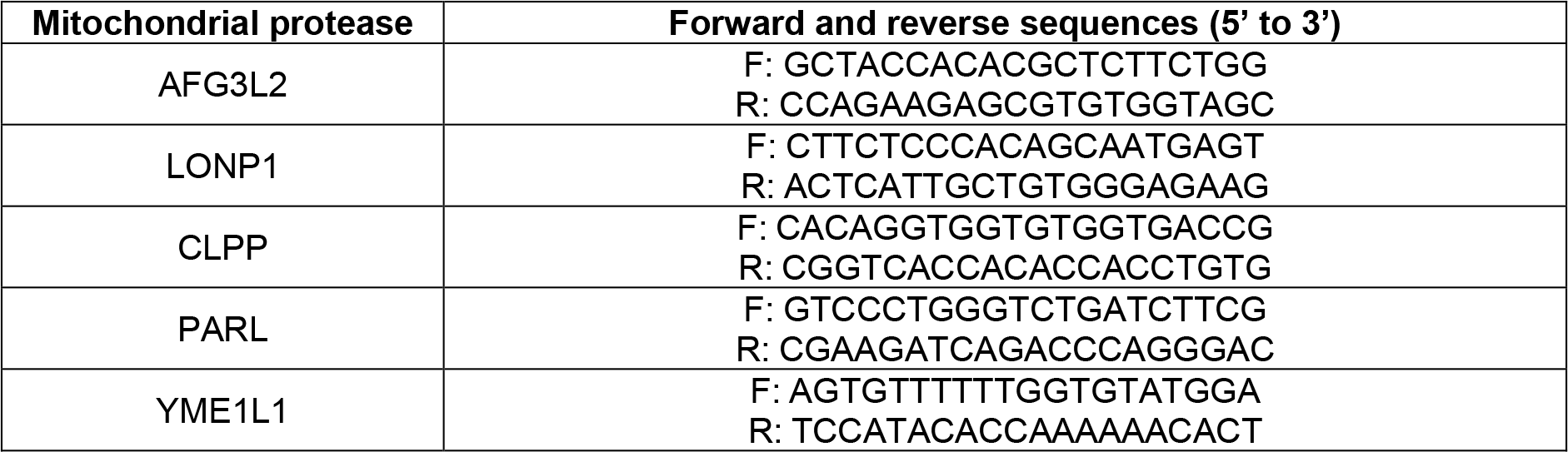
sgRNA sequences for mitochondrial proteases.

### CRISPR KO of mitochondrial proteases

Approximately 300,000 cells were seeded in a 6-well plate. After a 24-h recovery, the cells were transfected with 2 μg pX330-sgRNA and 0.2 μg of GFP empty vector using Lipofectamine 3000 (Invitrogen) at 1:3 DNA to lipofectamine following the manufacturer’s protocol. After 48 h of transfection, the cells were harvested and GFP+ cells were sorted into single cells in a 96-well plate using the BD FACSAria Cell Sorter. After 2–3 weeks, single-cell colonies were transferred to 6-well plates and genomic DNA was extracted after the cells reached ∼80% confluency. PCR was performed with the genomic DNA using forward and reverse primers located at least 150 bp from the sgRNA cut-site, yielding PCR amplicons from 400–800 bp. PCR products were purified and sent for Sanger sequencing. Sequencing results were analyzed with the Synthego Inference of CRISPR Edits (ICE) tool, and positive KO clones were further confirmed with next-generation sequencing.

### Analysis

For Western blots, densitometric analysis of the protein bands was performed using Image J. In CHX-chase experiments, relative protein abundance at different time points was normalized to the initial time point (0 hr). For immunofluorescence, 10 images totaling 200–400 cells were analyzed for the average FLAG and citrate synthase signals at each time point using Image J. All results were plotted as mean + standard deviation from the mean (S.D.M.). Statistical significance was determined using Student’s t-test, with * indicating *p* < 0.05 and ** indicating *p* < 0.01.

## ACKNOWLEDGEMENT

We thank Dr. Monther Abu-Remaileh (Stanford University) for the kind donation of the Atg5 KO HEK293T cells. We thank all members of the Kory lab for helpful discussions and Dr. Julie Gosse for editing the manuscript. This work was supported by an R35/MIRA (GM151097, N.K.) and an A*STAR fellowship (S.T.).

## AUTHOR CONTRIBUTIONS

R.Z.D. and A.D. generated the pX330-sgRNAs constructs and the CRISPR KO cells of mitochondrial proteases. A.D. and S.T. performed and analyzed experiments. N.K. and S.T. conceptualized the study and wrote the manuscript.

