## Supplementary Figures for "The iAAA-mitochondrial protease YME1L1 regulates the degradation of the short-lived mitochondrial transporter SLC25A38"

**Supplementary Fig. 1**

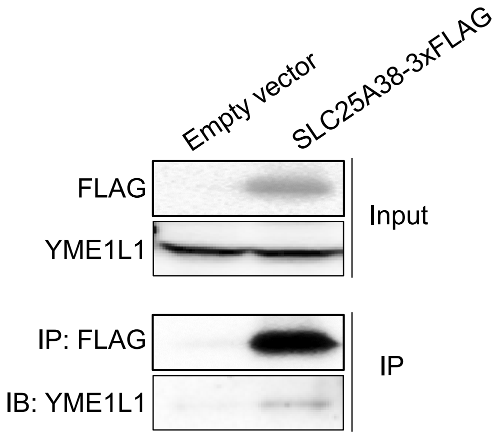


**Figure S1: Co-immunoprecipitation shows that YME1L1 interacts with SLC25A38 under basal condition.** Immunoprecipitation of FLAG in HEK293T cells stably expressing the empty vector and SLC25A38-3xFLAG followed by immunoblotting for YME1L1.

### Supplementary Fig. 1

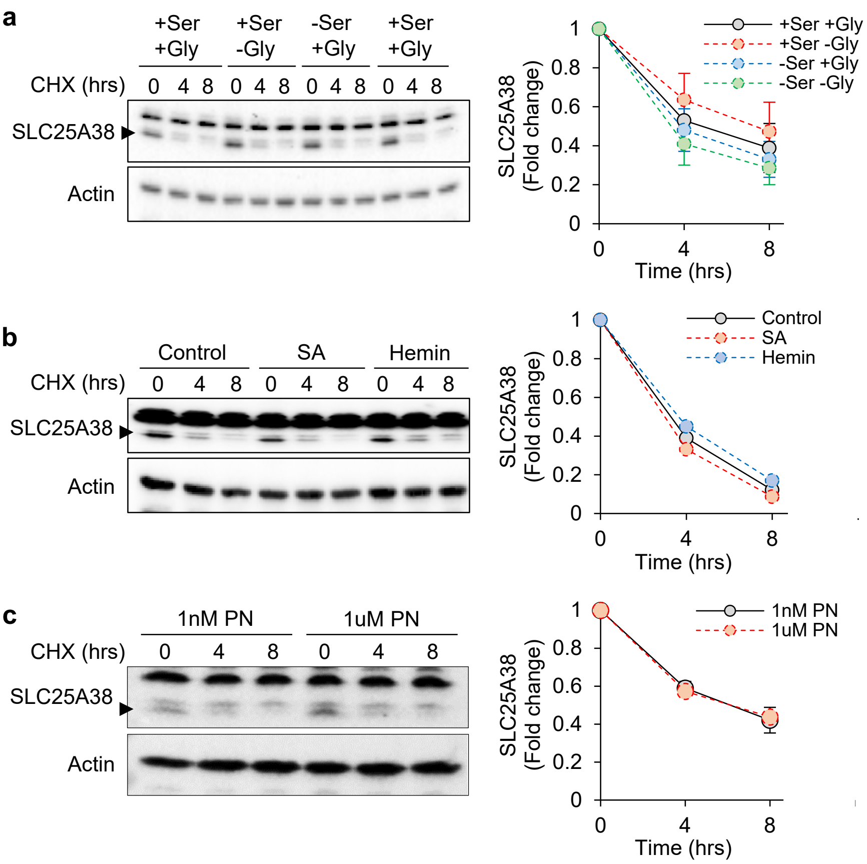


**Figure S2: Depletion of serine and glycine in culture media and inhibition of heme biosynthesis did not affect SLC25A38 stability.** (a) HEK293T cells were conditioned in serine and glycine depletion media, or depletion media supplemented with 0.4mM serine and/or 0.4mM glycine, for 24 hrs. The next day, 10ug/ml CHX was added and cells were harvested at different time points (n = 3). (b) HEK293T were conditioned in regular growth media (control), or media supplemented 0.5mM succinylacetone (SA) or 25uM hemin for 24 hrs. After 24 hrs, 10ug/ml CHX was added and cells were harvested at different time points (n = 1). (c) HEK293T were conditioned in vitamin B6 depletion media supplemented with 1nM (low) or 1uM (high) pyridoxine (PN) for 72 hrs. After 72 hrs, 10ug/ml CHX was added and cells were harvested at different time points (n = 2). Graphs show relative SLC25A38 levels at different CHX treatment time points under the different conditions. Fold change values were normalized to the protein abundance at 0 hrs. Values are mean ± S.D.M.
